# A novel method to selectively elicit cold sensations without touch

**DOI:** 10.1101/2022.06.14.496120

**Authors:** Ivan Ezquerra-Romano, Maansib Chowdhury, Caterina Maria Leone, Gian Domenico Iannetti, Patrick Haggard

## Abstract

**Background:** Thermal and tactile stimuli are transduced by different receptor classes. However, mechano- and thermo-sensitive afferents interact at spinal and supraspinal levels. Yet, most studies on responses to cooling stimuli are confounded by mechanical contact, making these interactions difficult to isolate. Methods for precise control of non-mechanical thermal stimulations remain challenging, particularly in the cold range.

**New Method:** We developed a non-tactile, focal, temperature-controlled, multi-purpose cooling stimulator. This method controls the exposure of a target skin region to a dry-ice source. Using a thermal camera to monitor skin temperature, and adjusting the source-skin distance accordingly, we could deliver non-tactile cooling stimuli with customisable profiles, for studying different aspects of cold sensation.

**Results:** To validate our method, we measured absolute and relative thresholds for cold sensation without mechanical contact in 13 human volunteer participants, using the method of limits. We found that the absolute cold detection threshold was 32.71°C ± 0.88 °C. This corresponded to a threshold relative to each participant’s baseline skin temperature of -1.08 °C ± 0.37 °C.

**Comparisons with Existing Method:** Our method allows cooling stimulation without the confound of mechanical contact, in a controllable and focal manner.

**Conclusions:** We report a non-contact cooling stimulator and accompanying control system. We used this to measure cold thresholds in the absence of confounding touch. Our method enables more targeted studies of both cold sensory pathways, and of cold-touch interactions.

**Highlights:** Most studies on cold sensation fail to control for concomitant tactile input.

A method to deliver non-tactile cooling stimuli was developed.

The method combines dry ice, a thermal camera, and motorised stages.

The method delivers rapid ramps and feedback-controlled pulses.

Thresholds for contactless cold perception were estimated in humans.

## 1 Introduction

The first step of thermoception is the activation of free nerve endings in the epidermis. However, contact thermal stimuli unavoidably coactivate deeper, touch-related afferents. These tactile signals interfere with thermonociceptive input both at spinal and supraspinal levels (Cahusac & Noyce, 2007; Ho et al., 2011; Mancini et al., 2015). As a result, both physiological and psychophysical responses to Aδ cooling-responsive units are confounded with tactile inputs, and quite possibly modulated by them. Therefore, cooling-mechanical co-stimulation precludes the study of non-tactile cooling responses. Yet, most research on cooling responses has used mechanical contact stimulators (e.g. Duclaux et al., 1974; Green, 2009). The logical way to study these interactions would involve comparing the effects of a cooling stimulus with the effects of a combined cooling and mechanical stimulus.

Therefore, studies of cold sensation would benefit from a cooling stimulation technique, which does not involve skin contact and mechanoreceptor activation such as laser stimulation in research on warm sensations (Iannetti et al., 2004). This technique would in turn allow interactions between cold and other sensations to be studied. Previous studies have attempted non-tactile cooling with stimulators that use ultrasound, chemicals, air flow or dry ice (CO_2_ solid form). These approaches have some advantages, but importantly have limitations for studying interactions between mechanical and cooling signals in the context of sensory binding and object perception.

A recent study presented a new method to cool the skin with a non-contact tactile display driven by ultrasound waves (Nakajima et al., 2021). While this method allows precise spatial and temporal control of the cooling stimulation, it does not selectively elicit cold sensations because ultrasounds generate vibrotactile sensations. In contrast, chemical approaches (e.g. menthol) specifically elicit cold sensations without mechanical pressure, but controlling the duration and intensity of chemically-induced sensations is limited (Typolt & Filingeri, 2020). Non-contact methods based on blowing chilled air at tissues allow precise control of the duration, area and intensity of stimuli, but involve a certain level of air pressure that presumably stimulate mechano-sensitive afferents (Murphy et al., 2001; Bujas, 1937). Radiation or convection methods might achieve cooling without activation of tactile afferents. However, studies which have achieved temperature decrease of the skin using radiation and/or passive convection transfer between dry ice and the skin lacked precise spatial and temporal control (Cataldo et al. 2016; Ferrè et al., 2018; Hardy & Oppel, 1938). Thus, they could not produce point-estimates of cold perceptual sensitivity of the kind used in psychophysical perceptual testing.

We have therefore developed a non-tactile, focal, temperature-controlled, multi-purpose cooling stimulator suitable for psychophysical testing on the perceptual aspect of cold sensation. Here, we describe three potential stimulation scenarios using this system. We validated our system in a study of human thresholds for cold perception in the absence of touch.

## 2 Materials and methods

To deliver non-tactile cooling stimulation, we used dry ice (CO_2_). The dry ice was held in a container, which varied in shape and dimension (10-2000 mL) according to the experimental application. The container was secured on a wooden support (30×2×2 cm), which could be moved in three axes using motorised linear stages (Fig. 1A) (A-LSQ150B and X-LSQ150B series, Zaber Technologies Inc.). To control the time of exposure of the skin to the dry ice, we placed a polystyrene shutter below the syringe tip, and controlled the shutter with a servo motor (SG90 Micro Servo Motor KY66-5, Longruner), driven by an Arduino Mega 2560. The shutter was closed between trials. To measure the temperature of the skin immediately below the syringe tip, we used a thermal camera (module temporal resolution: 8.7 Hz, Field of View: 57° & camera resolution: 60 × 120) (Lepton 3.5, Teledyne FLIR), interfaced with a computer through a I/O module (PureThermal 2 - FLIR Lepton I/O Smart module, Teledyn FLIR).

**Fig. 1.**
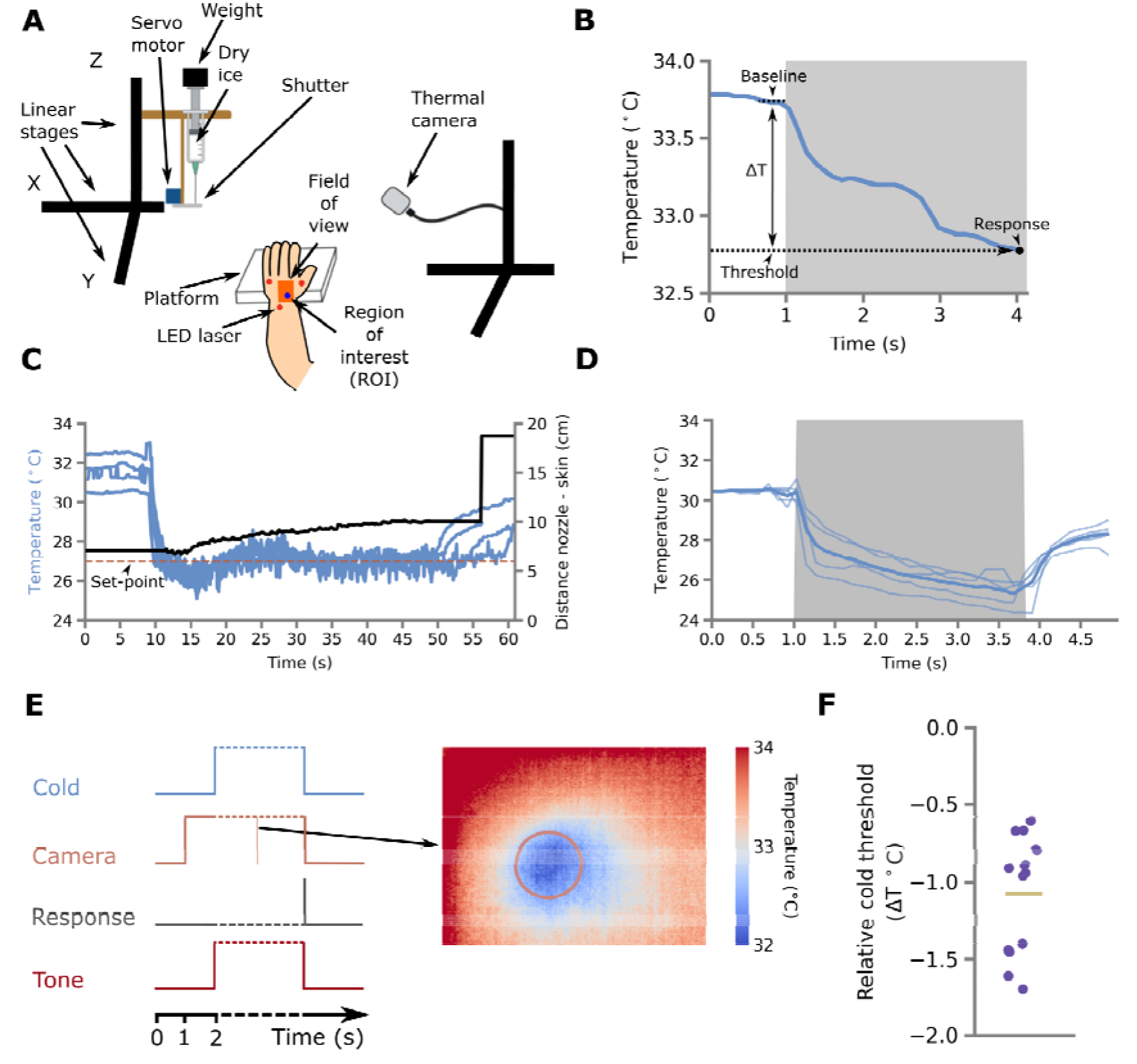
**A)** An illustration of the set-up with the main components. **B)** Example of focal cooling stimulation for determining cold thresholds by method of limits (scenario 1). The trace shows mean skin temperature in the skin ROI. The threshold level at which the participant first reported detecting cold is expressed relative to baseline (ΔT). The grey zone indicates the duration of cold stimulation (shutter is open). **C)** Example traces (blue lines, n=4 repetitions) of feedback-controlled stimulation (scenario 2). The horizontal, dashed red line indicates the set-point for the PID controller. The height of the stimulator above the skin is adjusted by PID control to achieve the desired temperature (one illustrative trace is shown by the black line, referring to right-hand ordinate scale). **D)** Examples of rapid, large-area cooling ramps (thin blue lines, n = 5 normalised repetitions; scenario 3). The thick blue line shows the mean. **E)** The panel to the left is a schematic displaying the temporal sequence of events in an MoL trial. The panel to the right shows a thermal image during dry ice stimulation. The orange circle represents the ROI. **F)** Cold detection thresholds relative to baseline skin temperature (ΔT) of 13 participants. Each datapoint represents the mean of 40 threshold estimates based on the method of Fig. 1B. The horizontal yellow line represents the mean.

To stimulate different skin regions during the same experiment, we had to control the position of the thermal camera. We used a second set of 3 motorised linear systems (2 stepper motor controllers LSM100B-T4 and 1 stepper motor controller LSM200B-T4, Zaber Technologies Inc.), which interfaced with the computer through controllers (2 stepper motor controllers X-MCB2 and 1 stepper motor controller X-MCB1, Zaber Technologies Inc.). These moved the thermal camera under computer control.

The motorised linear systems were positioned relative to the participant’s left hand. Three red lasers (5V 650nm 5mW, HiLetgo®) fixed to the wall pointed at the hand dorsum. The participant’s skin was marked with ink at the beam locations, so the experimenter could visually monitor the position of the participant’s hand throughout (Fig. 1A). To control the hardware, build the experiment and analyse the data, we wrote custom Python and Arduino code (see software repository: https://github.com/iezqrom/publication-cold-sensation-without-touch).

Using these general principles, we realised three separate stimulation scenarios suitable for psychophysical and neurophysiological experiments. Here, we do not report combining these scenarios, but in principle they could be combined to realise other scenarios. Firstly, we developed a focal cooling stimulus to measure cold thresholds relative to baseline. Secondly, we developed a temperature feedback-controlled Proportional-Integral-Derivative (PID) solution for delivering prolonged customised cooling profiles. Thirdly, we produced a wide area rapid-cooling thermal pulse, designed to investigate cooling-evoked EEG potential. Finally, to validate our method, we collected psychophysical data for scenario 1 and stimulation data from scenarios 2 and 3.

### 2.1 Scenario 1: Focal cooling stimulation for cold threshold measurements

In this scenario the dry ice was contained in a 10-ml syringe with a 4-cm blunt needle (BD Emerald Hypodermic Syringe - Luer Slip Concentric, BD). The syringe was wrapped in aluminium foil to reduce thermal loss. To obtain a constant pressure on the dry ice powder throughout, a weight of 1600 g was placed on the syringe plunger and a continuous rotation servo (FS5106R, Feetech) pressed on the plunger and the weight via a 3D printed linear system (Fig. 1A).

Thermal image recording started 2 s before the shutter opened. After shutter opening, the skin immediately below the syringe tip was exposed to convection of cooled air cooled by the dry ice (Appendix B). Further, the skin lost heat to the cooled syringe by radiation. As a result, the skin temperature gradually decreased, as recorded by the thermal camera. A region of interest (ROI, Fig. 1A & E) under the syringe tip was selected for online image analysis. Because the thermal camera was located slightly to the side of the syringe, and therefore had an oblique view, the circular ROI in the image had an elliptical projection (3.4×3.3 mm) on the skin. The pixel values in degrees Kelvin (K) were transformed into degrees Celsius (°C) and the temperature of the ROI was obtained by performing the mean across all the pixels within the ROI.

The capacity of the device to cool the skin decreases as the distance from the tip of the needle to the skin increases. We constructed a linear regression model (Appendix A) for selecting an appropriate distance to achieve the desired temperature range. For this scenario, we used a 5-cm distance.

To measure cold thresholds, we used the method of limits. At the start of each trial, a tone alerted the participant, and the shutter opened at the same time, exposing the participant’s skin to the dry ice, and leading to a progressive decrease of the measured temperature in the ROI. The participant was instructed to press a foot pedal when they first felt a cold sensation on the stimulated skin region. When the pedal was pressed, the stimulator shutter closed, the tone terminated, and the final skin temperature was stored (Fig. 1E). To allow skin temperature to return to baseline, 4 locations were randomly stimulated. The locations were arranged in a square grid with a spacing of 2 cm. The same location was restimulated only after 2 other locations had been visited, ensuring a minimum of 30 s for thermal recovery between stimulations at each site.

As well as measuring the absolute threshold, we used the data from each trial to calculate the relative threshold, i.e., the smallest drop in temperature from baseline that the participant could detect (ΔT) (Hafner et al., 2015). Unlike contact thermal stimulators, our stimulator does not set the initial temperature of the skin before each stimulation like the lasers used in the study of heat sensation (Iannetti et al., 2004). We observed variability (mean: 33.8 ± 0.9 °C SD) in the baseline skin temperature across participants and grid locations. Because we only restimulated one location after at least 30 s, this variability is probably unrelated to the experimental procedure. Thus, ΔT is arguably a more ecologically valid threshold measurement of cold sensitivity than an absolute measurement. Our method and procedure are suited to account for this variability.

To measure ΔT, an average of the baseline skin temperature within the ROI was taken across 26 frames in the 300 ms before shutter opening. Then, the temperature of the ROI upon pedal press was subtracted from the previous averaged baseline value to obtain ΔT (Fig. 1B).

#### Participants and ethics

We measured cold thresholds on the hand dorsum of 13 participants (mean age: 24.1 ± 3.5 SD), obtaining 40 estimates per participant. Appropriate risk management procedures, notably around handling dry ice, were implemented. The research protocol was approved by the UCL Research Ethics Committee (ID number: ICN-PH-PWB-0847/010). Room temperature fluctuations and air currents were minimised by closing windows and doors.

### 2.2 Scenario 2: Feedback temperature-controlled custom cooling stimulation

This scenario allowed delivery of temperature-controlled non-tactile cooling stimulation for psychophysical paradigms which require long periods of constant low temperature. Raising the cooling source resulted in less cooling, while lowering it towards the skin produced more cooling. Therefore, to achieve the desired constant temperature reading from the thermal camera, we continuously adjusted the distance between dry ice and skin (Fig. 1C).

A PID algorithm closed the feedback loop between thermal image and the cooling source height above the skin. First, a desired temperature is set. Then, the thermal camera detects the temperature of a skin ROI, and a simple PID feedback controller sends position commands to the motorised linear system, adjusting the height of the container until the desired temperature is reached. This allows precise temporal and spatial control of skin temperature for psychophysical experiments. For instance, in combination with a non-tactile warm stimulator, a temperature-controlled radiant Thermal Grill Illusion (TGI) could be elicited for the first time.

In this set-up, the dry ice was held in a carboard container. The cardboard container had dimensions 10.2×10.2×21.8 cm with a total volume of 1600 ml. It was filled with 300 g of dry ice. The base was perforated with a 6-mm diameter copper tube. For these studies, the ROI had a projected elliptical shape on the skin of 5×4 mm. The interior of the container was covered with foil and its exterior with polystyrene foam in order to limit thermal loss and convection to the copper tube.

### 2.3 Scenario 3: Rapid, wide-area, high-intensity cooling stimulation

This scenario allowed delivery of fast non-tactile cooling stimulation of a large skin area, designed to produce a strong afferent volley and an evoked brain response. Event-related EEG potentials require such strong stimuli with rapid onsets. Steep cooling ramps cause synchronous activation of many cold afferents, and therefore improve the signal-to-noise ratio of event-related EEG potentials, thus paralleling what has been demonstrated for steep radiant heating ramps (Iannetti et al., 2004).

In this set-up, the dry ice container for scenario 2 was used, but the base was perforated with three outlet tubes to increase the stimulation area. Therefore, the exposed skin area was larger, and the skin ROI was a 10×8 mm ellipse. Stimulation led to a rapid temperature decrease at 13 ± 3 °C/s SD (Fig. 1D). Previous studies have shown that a cooling ramp of 10-17°C/s delivered to an 1444 mm^2^ skin area is sufficient to detect a reproducible evoked potential (Duclaux et al., 1974). Thus, our stimulation method permits rapidly cooling a large patch of skin without touch, and potentially measuring the EEG responses elicited by cooling stimuli without mechanical input.

## 3 Results

Our stimulator successfully delivered cooling stimulation without touch. We determined absolute and relative temperature threshold of cold detection (Fig. 1B & F). We also achieved repeatable, sustained, temperature-controlled cooling stimulation suitable for psychophysical studies of cold perception with stimuli varying in intensity, location, and duration (Fig. 1C). Finally, we demonstrated a rapid decrease in skin temperature without touch, which is suitable for electrophysiological recordings (Fig. 1D).

Baseline temperature before cooling stimulation was 33.8 ± 0.9 °C SD across 13 participants. The absolute threshold for reporting cooling stimulation was 32.7 ± 0.9 °C SD across participants. Thus, the relative threshold for cold detection (ΔT) was - 1.1 ± 0.4 °C across participants (Fig. 1F).

## 4 Discussion

We report a novel method to generate controlled and focal cooling stimulations without the confounds of mechanical input. We show how the system can be used to perform psychophysical experiments on the perceptual aspect of cold sensation. We have focussed on the methodological issues around non-contact cooling stimulation. Importantly, our approach overcomes some of the limitations of previous non-contact cooling approaches. Firstly, non-contact methods using ultrasound and air blow do involve mechanical stimulation (Bujas, 1937; Nakajima et al., 2021; Murphy et al., 2001). Secondly, previous studies using chemical and dry ice approaches to selectively elicit cold sensations had poor spatio-temporal controllability (Cataldo et al. 2016; Ferrè et al., 2018; Hardy & Oppel, 1938; Typolt & Filingeri, 2020).

Furthermore, we show that our method can be used for perceptual psychophysics. The most widespread cold perception test is the calculation of a cooling detection threshold using a method of limits. This is the basis of the cold threshold test performed in Quantitative Sensory Testing (QST) studies that are frequently used in clinical studies. The data from our device (Fig. 1F) showed cold thresholds close to the published normative values for QST using methods of limits (Rolke et al., 2006: 30.9°C from 32°C baseline; Hafner et al., 2015: -1.0°C degrees). All these studies, however, used contact thermodes, thus introducing a tactile element. Here, we have shown that our method could reliably estimate focal cold detection thresholds without mechanical stimulation, which are comparable to existing normative values. This validates the use of our method for perceptual psychophysical experiments. Crucially, the duration and size of our cooling stimuli is unlikely to trigger homeostatic responses. Thus, our method is suitable for studying the perceptual aspect of cold sensation, but not its regulatory aspect.

Therefore, our non-tactile thermal cooling stimulator opens the possibility of investigating cold-touch interactions. The classic method to study such interactions would involve comparing the responses to thermal stimuli both with and without concomitant mechanoreceptor stimulation. However, the mechanisms of thermo-tactile interactions are poorly understood because most methods for delivering cooling stimulation involve mechanical stimulation. In future studies, our stimulator will be used to address the scientific question of how mechanical stimulation interacts with cooling stimulation. For example, cold thresholds could be measured in the presence or absence of concomitant touch. If an interaction between touch and thermal sensitivity is established, EEG studies could investigate the neurophysiological mechanisms underlying this interaction. Our rapid, large-scale cold stimulator could potentially allow future electrophysiological recordings of cold-evoked potentials, and of how they might be modulated by tactile input. Furthermore, future studies could combine scenarios to deliver more sophisticated thermal profiles such as PID-controlled ramps (scenarios 2 & 3). This combined scenario could be used to study cold perception for different temperature gradients without tactile input.

Our results of the cooling detection thresholds have some limitations. First, the method of limits does not distinguish between two key components of sensory detection: sensitivity and bias. Further, we did not include ‘catch’ trials, in which the auditory tone would occur without any cold stimulation. The absence of catch trials could potentially induce a response bias, with participants responding based on the expectation that a cold sensation would occur. Thus, our measures of cold thresholds should not be taken as perfect estimates of sensitivity. However, this does not detract from the scientific value of the stimulator apparatus. Future studies could use psychophysical methods such as signal detection to estimate thermal sensitivity independent of bias.

In our method, two modes of heat transfer contribute to skin cooling: convection and radiation. Convection cooling takes place as sublimated CO_2_ and cooled air flow down from the container to the skin because they are more dense than ambient air. Radiative cooling also transfers thermal energy from warmer objects (the skin) to cooler ones (the stimulator). Our design cannot distinguish the respective contributions of convection and radiation to skin cooling. The very focal cooling achieved in scenario 1 suggests that convection dominates. One might object that cold air currents flowing downwards to the skin constitute a mechanical stimulus, and that our method is not therefore completely non-tactile. We addressed this limitation by measuring the convection airflow in our exposure Scenario 1 with a Pitot tube, and calculating the resulting mechanical forces at the skin. The velocity of the airflow immediately below the syringe tip was measured as 4.0 m/s ± 0.30 SD (density of sublimated CO_2_: 1.836 kg/m^3^; MPXV7002DP pressure sensor, NXP). Calculations confirmed that the resulting forces on the skin were below published mechanoreceptor threshold values. Finally, in informal pilot testing, we gently blew air through the syringe at this velocity, and found that this level of airflow was not perceptually detectable (Appendix B). Therefore, our cooling stimulator could be considered having both high sensitivity (effectively stimulating cold afferents) and high specificity (not stimulating non-thermal afferents, notably mechanoreceptor afferents). However, further psychophysical and electrophysiological studies should investigate the sensitivity and specificity of our method.

## 5 Conclusions

We describe a novel non-tactile, focal, temperature-controlled, multi-purpose stimulator. We show how this device can be used for non-tactile thermal stimulation in humans. Future studies can use our stimulation method in different psychophysical and neurophysiological experiments to establish a thermo-tactile interaction. Understanding the mechanisms of thermo-tactile and thermo-thermal interactions would help improving clinical treatments, thermal displays and other haptic devices.

## Authors’ contributions

Ivan Ezquerra-Romano: Conceptualization, Methodology, Software, Data collection, Data curation, Visualization, Investigation, Writing - Original, Validation, Formal analysis, Investigation. Maansib Chowdhury: Data collection, Validation. Caterina Maria Leone: Conceptualization, Methodology, Draft, Writing – Review & Editing. Gian Domenico Iannetti: Conceptualization, Methodology, Funding, Supervision, Resources, Draft, Writing – Review & Editing. Patrick Haggard: Conceptualization, Methodology, Funding, Supervision, Writing – Review & Editing, Resources, investigation.

## Acknowledgements

I.E.R. was supported by the Biotechnology and Biological Sciences Research Council (UK) [grant number BB/M009513/1]. M.C. was supported by a European Union Horizon 2020 Research and Innovation 385 Programme (TOUCHLESS, project No. 101017746). Additional funding was provided by a UCL ‘Cities Partnership Project’ grant, which supported C.M.L. and G.D.I. G.D.I. was supported by the ERC (PAINSTRAT grant). The authors would like to thank Martin Donovan for his technical support in the development of the methodology. Help from Dr. Shinya Takamuku was also greatly appreciated. Informal discussions with Angel Ezquerra and Guanhaven Romano-Mendoza were helpful in the inception and development of the methodology.

## Appendix A

**Linear regression model**

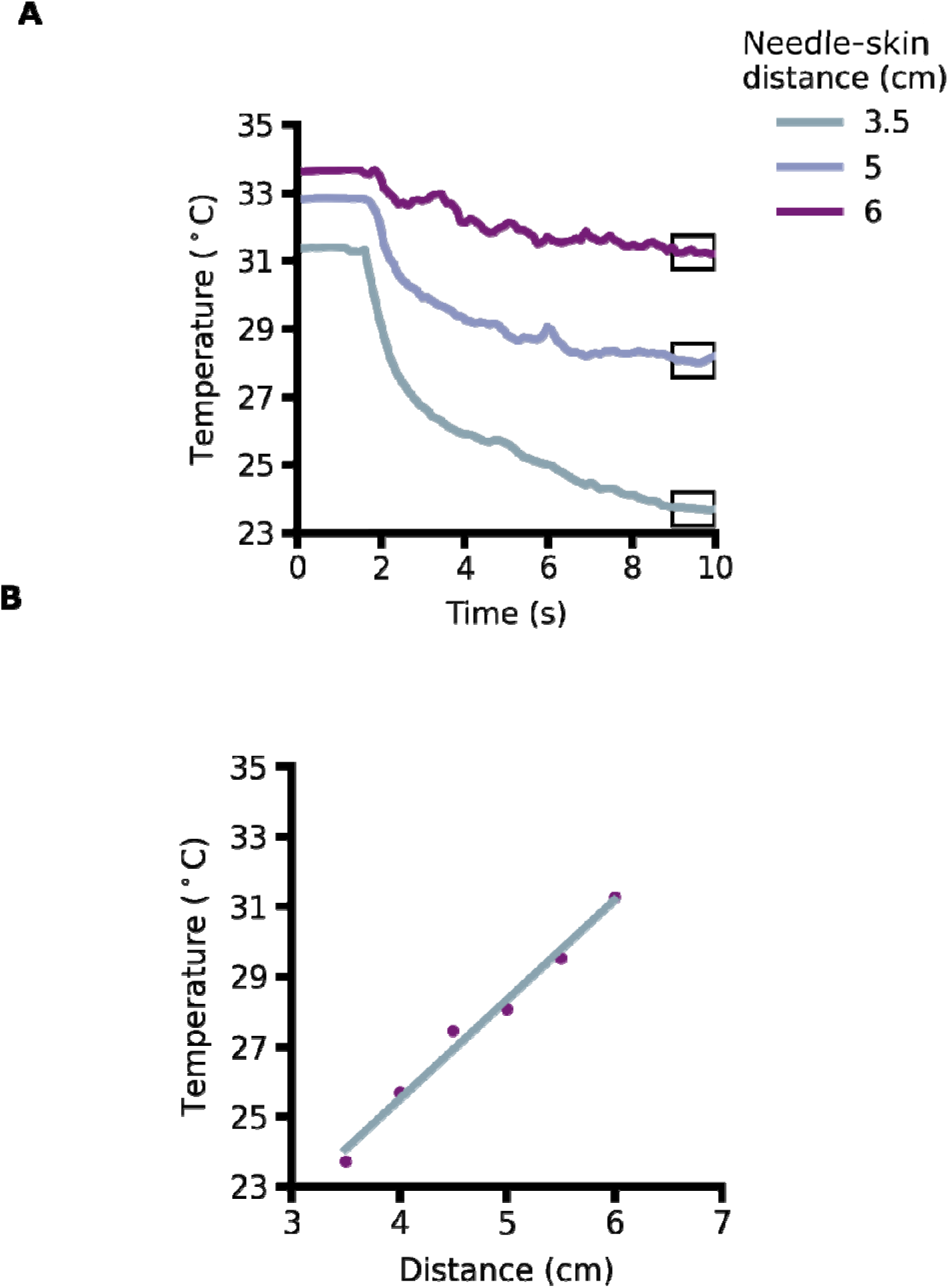
Linear regression model. Thermal effects of distance between cold source and skin are used to create a linear regression model for obtaining feedforward stimulus parameters. A) Traces showing mean temperature of the skin ROI for 3 different distances between the needle’s tip and the skin. Each trace is the mean of 5 recordings at the given distance. The grey zone indicates the duration of cold stimulation (shutter is open). The black open boxes over indicate the data used to calculate the model represented in Fig. B. B) Dots showing the final temperature at each distance and trace showing the linear fit. At 6 different distances, 5 recordings of the skin temperature during cold stimulation were made. After performing the mean of the traces at each distance, the mean temperature during the final 1 s of stimulation was extracted (black box in Fig. A). A linear regression was used to model the relation between distance and temperature. The model could then be used to position the syringe either to allow a desired temperature to be reached, either using feedforward control, or as an initial estimate of distance for a feedback-controlled stimulation.

## Appendix B.

**Airflow force and d’ calculations.**

### Airflow force calculation

Dry ice is solidified CO_2_. It sublimates directly from its solid state below -80°C to a gaseous state at standard temperature and pressure. These characteristics make this material suitable for non-tactile cooling. The sublimation of dry ice produces airflow from the needle’s tip of the syringe. To obtain the force that this airflow exerts on the skin, we use the following formula for fluid dynamics:

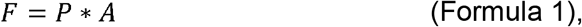

To calculate the force, we need to obtain the pressure (P) and the area (A) of the airflow when it collides with the skin.

In a fluid, the dynamic pressure is the kinetic energy per unit volume. To calculate the dynamic pressure, we can use the following formula:

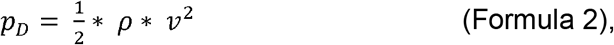

where *ρ* is the density of the fluid and *ν* is the speed of the fluid. We used the density of CO_2_ at standard room temperature (25 °C) and pressure (1 atm) - *ρ* = 1.84 kg/m^3^. The velocity of the jet of air was measured with a pitot tube. The pitot tube was placed at 5 cm below the tip of the needle, which is the distance at which the skin was during the experiment described in Fig. 1F. The mean velocity of the jet of air sampled at 10 Hz and averaged over a 4 s period was 4.06 m/s (SD 0.30 m/s). Therefore, following from Formula 1, the dynamic pressure that the jet of air exerts on the skin is:

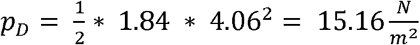

To obtain the area of the airflow when it collides with the skin, we can use the ellipse drawn on the skin by the circular ROI taken from thermal camera measurements in scenario 1. The cooled area of the skin was measured as an ellipse with axis lengths 3.37 mm and 3.32 mm. The area of an ellipse is:

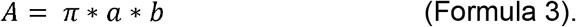

Therefore, the area cooled in scenario 1 was,

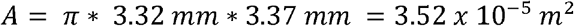

Following from Formula 1, the force that the jet of air exerts on the skin is:

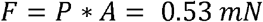

The estimated threshold for exciting a single mechanoreceptor afferent by punctate stimulation of glabrous skin was estimated using microneurography (Johansson et al., 1980). They found that RA units had a median threshold of 0.58 mN. PC units had a median threshold of 0.54 mN. Slowly adapting SAI and SAII units had median values of 1.3 mN and 7.5 mN, respectively. In our setup, convection currents from the cold source may be assumed constant. Therefore, the mechanoreceptor afferents most likely to be stimulated are the SAI units. Therefore, the mechanical element of our cold stimulation is less than half the force level suggested to trigger a single mechanoreceptor afferent action potential. We therefore conclude that convection currents from our cold stimulator were unlikely to produce any effective mechanical stimulation.

### D’ for the detection of airflow

To further assess whether the mechanical stimulation generated by the minimal airflow during our main experiment was perceptually detectable, we performed a pilot Signal Detection Theory experiment.

The experimenter blew through the syringe on a participant’s forearm. In 10 trials, the syringe was perpendicular to the skin and at distance of 5 cm (stimulus present). In another 10 trials, the syringe was moved away so that the participant’s arm was not stimulated (stimulus absent). The participant was asked to detect the jet of air. For this experiment, 4 naïve, blindfolded participants were tested.

Before performing the experiment, the experimenter was trained to blow through the syringe to generate a jet of air with a velocity of 5.09 m/s as recorded by the Pitot tube. Therefore, the airflow of atmospheric air (density: 1.204 kg/m^3^) produced a force of 0.55 mN.

On average, the hit rate was 26% and the false alarm rate was 15%. Across 4 participants, D’ was 0.53 and the criterion response (c) was -0.94. This pilot psychophysical test suggests that the level of airflow (speed and area) generated by the syringe with dry ice was below perceptual detection threshold for mechanoreceptor sensations.

